# Postural effects on arm movement variability are idiosyncratic and feedback-dependent

**DOI:** 10.1101/2020.12.23.424234

**Authors:** Preyaporn Phataraphruk, Qasim Rahman, Kishor Lakshminarayanan, Mitchell Fruchtman, Christopher A. Buneo

## Abstract

Reaching movements are subject to noise arising during the sensing, planning and execution phases of movement production, which contributes to movement variability. When vision of the moving hand is available, reaching variability appears to be strongly influenced by noise occurring during the specification and/or online updating of movement plans in visual coordinates. In contrast, when vision of the hand is unavailable, variability appears more dependent upon hand movement direction, suggesting a greater influence of execution noise. Given that execution noise acts in part at the muscular level, we hypothesized that reaching variability should depend not only on movement direction but initial arm posture as well. Moreover, given that the effects of execution noise are more apparent when movements are performed without vision of the hand, we reasoned that postural effects would be more evident when visual feedback was withheld. To test these hypotheses, subjects planned memory-guided reaching movements to three frontal plane targets, using either an “adducted” or “abducted” initial arm posture. Movements were then executed with and without hand vision. We found that the effects of initial arm posture on movement variability were idiosyncratic in both visual feedback conditions. In addition, without visual feedback, posture-dependent differences in variability varied with movement extent, growing abruptly larger in magnitude during the terminal phases of movement, and were moderately correlated with differences in mean endpoint positions. The results emphasize the role of factors other than noise (i.e. biomechanics and suboptimal sensorimotor integration) in constraining patterns of movement variability in 3D space.

## Introduction

Variability is inherent in movement production and studies of movement variability have and continue to inform our understanding of coordinate transformations, motor learning, and optimal motor control (Gordon et al., 1994;McIntyre et al., 1997;Harris and Wolpert, 1998;McIntyre et al., 1998;van Beers, 2009). Movement variability has been attributed in part to “neural noise” arising during the encoding and integration of sensory signals and/or the planning and generation of motor commands (Faisal et al., 2008). Until fairly recently conventional wisdom has held that noise is detrimental to motor behavior (Harris and Wolpert, 1998;van Beers et al., 2002;Herzfeld and Shadmehr, 2014). However, recent work has shown that a component of movement variability appears to arise from a gradual accumulation of the random effects of planning noise, a phenomenon which could benefit motor learning by fostering exploration in the motor command space (van Beers et al., 2013). Similarly, the observation that baseline levels of movement variability can predict the rate at which individual human subjects learn motor tasks suggests that some component of neural noise might actually be advantageous or even necessary for motor learning to occur (Wu et al., 2014). Recent work demonstrating that a covariation of slow drifts in neural and behavioral variability is well explained by a simple model of error-corrective learning, appear to provide additional support for this idea (Chaisanguanthum et al., 2014).

For reaching movements, noise can arise during the initial encoding and/or updating of the hand and/or goal location (“sensory noise”), during the transformation of sensory signals into motor commands (“planning noise”) or during the transformation of commands into movement (“execution noise”) (Buneo et al., 1995;van Beers et al., 2004;Osborne et al., 2005;Churchland et al., 2006a;Churchland et al., 2006b;Shi and Buneo, 2012). As a result, the effects of noise on reaching variability are highly context-dependent. For example, when the hand is visible during movement, variability tends to be greater in depth than along other axes, reflecting the reduced precision associated with visual localization of the hand and/or targets in depth (McIntyre et al., 1997;1998;Carrozzo et al., 1999). In contrast, without visual feedback of the moving hand variability is greatest along an axis that is collinear with the direction of movement (Gordon et al., 1994;McIntyre et al., 1997;1998). These effects do not appear to be related to planning noise but noise associated with execution, particularly during the terminal phases of movement (van Beers et al., 2004;Apker and Buneo, 2012).

Other studies suggest that the effects of noise on variability should depend not only on movement direction but on arm posture as well. In simulation, patterns of movement variability induced by the introduction of planning and execution noise differed markedly when movements were initiated from different locations in a 2D workspace (therefore employing different arm postures), even for movements of the same planned direction and extent in Cartesian coordinates (Fig. 1). In other words, the effects of noise on variability were both movement direction- and posture-dependent. Studies of memory-guided reaches performed with diminished or absent visual feedback support this view. For example, when subjects were required to reach to memorized target locations from the same starting position using either the left or right hand, the spatial characteristics of movement variability depended upon both the starting position and the hand used to make the movement (McIntyre et al., 1998). Although effects of hand used were interpreted as effects of arm posture in this study, this interpretation was confounded by another factor: arm dominance. In other words, differences in variability for movements made from the same starting position but using different hands may have arisen at least in part from the different specialized functions of the dominant and non-dominant hemispheres (Sainburg, 2014), rather than due to differences in limb configuration per se.

**Figure 1.**
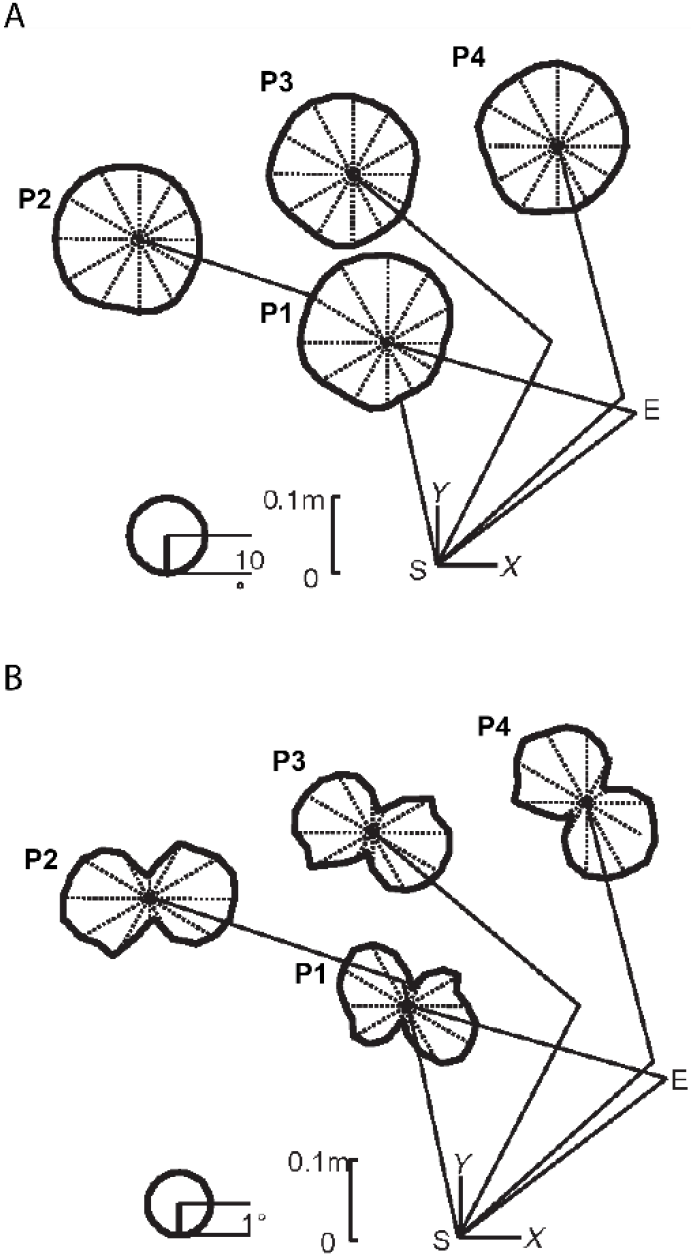
Results of feedforward, forward dynamics simulations examining the effects of planning noise (A) and execution noise (B) on arm movement variability in the horizontal plane. In both A and B, simulation results for four starting positions (S1-S4) are shown. Polar plots surrounding each starting endpoint position indicate the circular standard deviations of the directional errors resulting from the imposed noise for each of 24 movement directions. S: shoulder, E: elbow. Adapted from Shi and Buneo (2012).

In the present study, we tested the hypothesis that reaching variability was arm posture-dependent while controlling for effects of arm dominance and differences in planned endpoint trajectories. Memory-guided reaching movements were made with the right arm from a single starting position to three targets contained in a frontal pane. At the starting position, subjects matched one of two desired arm configurations (“adducted” and “abducted”) by rotating the arm about the shoulder-hand axis, thereby maintaining a constant endpoint position. In addition, movements were performed with and without vision of the moving hand. Based on previous studies we hypothesized that patterns of reaching variability would vary systematically with both movement direction and arm posture and that these variations would be more evident when vision of the moving hand is withheld.

## Methods

### Subjects

Ten subjects (5 men and 5 women) between the ages of 19 and 53 were recruited to perform the experiment. Subjects were briefed on the experimental procedure, which involved reaching to targets in 3D space using the same starting fingertip position but different initial arm postures, but were naïve to the actual purpose of the experiment. The protocol was approved by the Arizona State University Institutional Review Board (IRB) and subjects read and signed an IRB approved informed consent form before participating in the experiment, which was conducted in accordance with the principles expressed in the Declaration of Helsinki.

### Apparatus and Data Acquisition

The experimental apparatus consisted of a large, standing metal frame that supported a 3D stereoscopic monitor (Dimension Technologies, Rochester). The monitor projected images through an opening in the metal frame onto a reflective mirror embedded in a metal shield. The metal shield was oriented at a 45° angle with respect to the monitor and enabled the subjects to see the projected images on the mirror. The metal shield also served to block the subjects’ arms from view. Subjects positioned their heads on a chin rest which aligned their eyes with the center of the mirror and were asked not to look away from the mirror during the entire experiment. Subjects were also asked to limit repositioning of their body during the experiment.

An active-LED-based motion tracking system was used to track movements of the arm (Visualeyez VZ-3000 motion tracker; Phoenix Technologies, Burnaby, British Columbia; 250-Hz sampling rate; 0.5-mm spatial resolution). Three (3) LEDs were placed on each subjects’ fingertip, elbow and shoulder respectively. The position of the fingertip LED was fed back to the subject in near real-time within a virtual reality environment developed in Vizard (WorldViz, Santa Barbara, CA). The fingertip position, starting position and targets were displayed as green spheres of ~5 cm diameter in the VR environment. To aid in depth perception, a cube object was also rendered in the VR environment. Monitoring of fingertip position and arm configuration, as well as interfacing with the VR environment was accomplished via a custom program developed in LabVIEW (National Instruments Corporation, Austin, Texas) which also downsampled the fingertip position at 125 Hz.

### Experimental Design

Subjects were required to make memory-guided reaching movements to three targets using one of two initial arm configurations and with or without visual feedback of the fingertip. We used a memory-guided task to be consistent with previous studies of reaching in three dimensions (e.g. McIntyre et al. 1998). The starting position of the hand was located on the body midline at approximately shoulder level and all targets were located 11.7 cm from the starting position. One target was located directly ahead of the starting position on the body midline (target M) at an elevation angle of 60° with respect to the horizontal plane containing the starting position. The other two targets were located 45° to the left and right of the starting position (targets L and R, respectively) and at elevation angles of 45°. Targets L and R appeared at approximately eye level, while target M appeared slightly superior to the lateral targets. Given the arrangement of the targets, on a given trial subjects were required to reach upward and in depth from the starting position and either directly forward (0°) or slightly leftward (−45°) or slightly rightward (45°).

Trials began with the illumination of the starting position. Once subjects acquired the starting position with their fingertip and maintained that position for 1000 ms the start position was extinguished and a target was illuminated for 300 ms, which was then also extinguished. Subjects were then required to withhold making a movement to the target during an ensuing memory period of 500-1500 ms. At the end of the delay period a 60 Hz tone was generated, which served as the ‘go’ cue to begin the movement. If subjects completed their movements within 1000 ms and maintained their fingertip at the perceived target location for 1250 ms, another auditory tone was generated, indicating the end of the trial. Vision of the fingertip was available to the subject throughout the trial in the vision (V) condition but was removed at the go cue in the non-vision (NV) condition. Feedback condition (V, NV) and target direction were randomly varied on trial-by-trial basis.

Trials were organized into two blocks, with each block employing either an “adducted” or “abducted” arm posture at the starting position (Fig. 2A). Arm posture was changed by rotating the arm about the shoulder-hand axis. In this way, the same starting position was maintained, thereby ensuring that planned movement vectors were largely identical between postures. The order of the blocks were randomized across subjects. The angle that the arm plane (i.e. the plane containing the upper arm and forearm) made with horizontal was used to define arm posture. An arm plane angle of 0° corresponded to full abduction and 90° indicated full adduction. For the abducted block, subjects were required to maintain their posture between 0 and 45° and for the adducted block, a posture between 45 and 90° was required. If at any point during the trial subjects failed to maintain their posture within the required range, a 1kHz tone was generated, cueing them to reposition. A total of 90 trials were performed in each block (15 V trials and 15 NV trials to each of three targets). Subjects were given an approximately one minute rest period every 15 trials within a block as well as between blocks to minimize fatigue due to elevating the limb for extended periods.

**Figure 2.**
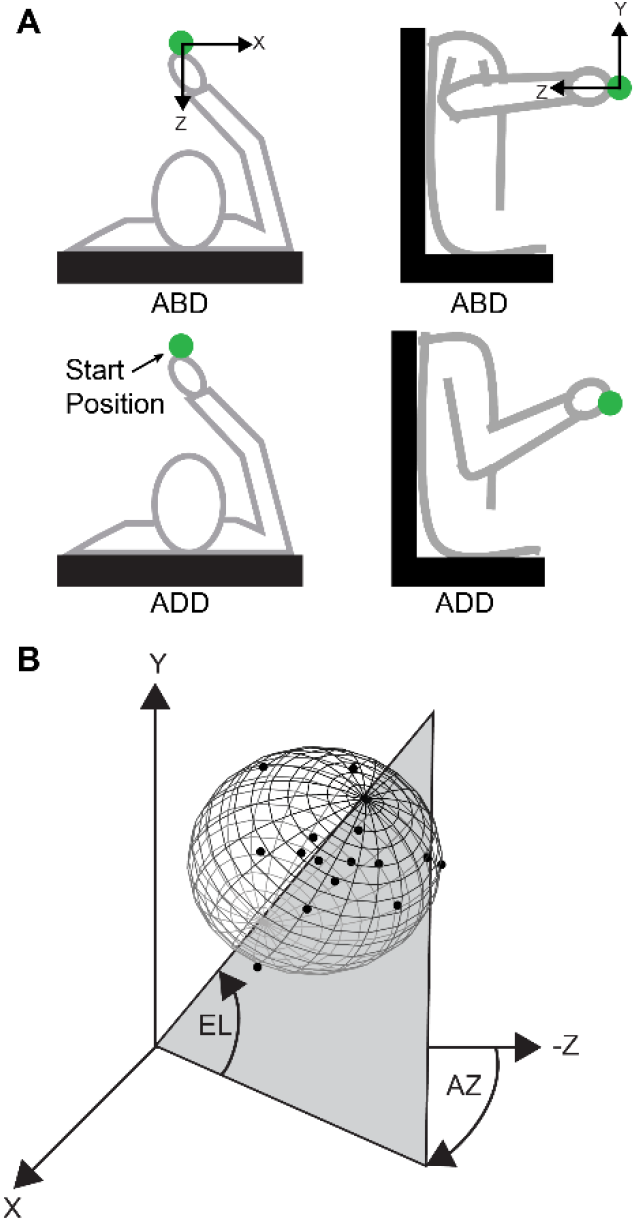
Arm postures and angles used to define variability ellipsoid orientations. A. Top-down (left) and lateral (right) views of the abducted (ABD) and adducted (ADD) postures. B. Simulated 3D endpoint distribution showing definitions for azimuth (AZ) and elevation (EL). Azimuth was defined as the angle in the x-z plane measured from the negative z-axis, with positive values being counterclockwise. Elevation was defined as the elevation angle from the x-z plane.

Subjects had no knowledge of the trial parameters and were given instructions to move quickly and accurately to the targets using a single uncorrected movement. A trial was considered successful if the subject maintained the initial starting position and arm posture, reached a target within the required spatial and temporal windows, and maintained position at the end of the movement for 1200 ms. If any of the criteria for a successful trial were not met, the trial was aborted and repeated later in the block.

### Data Analysis

Movement data were smoothed using digital low-pass filters (4^th^ order, 6 Hz cutoff). The beginning and end of each movement was defined as the points at which the tangential movement velocity exceeded or fell to 10% of its peak value. Data from trials where the tangential velocity exhibited multiple peaks or other irregularities were discarded (2% of all trials).

Movement endpoint data, sorted according to target direction, visual feedback condition, and arm posture, were characterized by their overall volume, aspect ratio, and orientation. To determine the volume and aspect ratio we first calculated 95% tolerance ellipsoids for each endpoint distribution (Khachiyan and Todd, 1993;Khachiyan, 1996;Moshtagh, 2009). The volume of each ellipsoid was then quantified as follows:

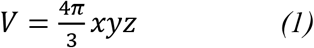

where x represents the radius of the major axis of the ellipsoid and y and z refer to the radii of the minor axes. We calculated the aspect ratio of each ellipse as the ratio of the radius of the major axis to that of the smaller of the radii of the minor axes. To quantify the orientation of each endpoint distribution, principal components analysis (PCA) was used (McIntyre et al., 1997;1998;Carrozzo et al., 1999;Apker et al., 2010;Apker and Buneo, 2012). The first eigenvector derived from PCA was used as an indicator of the principle axis of movement variability. The orientation of this axis was parameterized by its azimuth (angle in the x-x plane) and elevation (angle out of the x-z plane) (Fig. 2C).

### Statistical Analyses

For all analyses, an alpha of 0.05 was used.

#### Analyses within visual conditions

Visual conditions were initially analyzed separately for several reasons. First, the initial focus of this investigation was on effects of direction and configuration, not effects of vision, which we and others have previously analyzed extensively in similar tasks (e.g. Apker et al., 2010;2012). In addition, the previous studies which are most relevant to the present one also analyzed visual conditions separately (McIntyre et al., 1997;1998); thus to facilitate comparison with these studies we adopted a similar approach. Lastly, to our knowledge three or more factor analyses of variance for angular data do not exist. Therefore, for each visual condition we used two-factor ANOVAs to analyze both our angular (orientation) and non-angular data (volume, aspect ratio). More specifically, effects of movement direction and initial arm posture on ellipsoid volume and aspect ratio were analyzed using two-factor repeated measures ANOVAs implemented in SPSS version 25 (IBM Corp.). Analyses of ellipsoid orientation (azimuth, elevation) were conducted in Matlab R2019a (The Mathworks, Inc.) using the CircStat toolbox (Berens, 2009). Here, the Rayleigh test was used first to probe the distributions of azimuth and elevation angles for deviations from non-uniformity. Subsequently, effects of movement direction and initial arm posture on ellipsoid azimuth and elevation were analyzed using two-factor circular ANOVAs.

For each visual condition we also analyzed the within-subject differences in ellipsoid orientation between postures. For each visual condition, a linear regression analysis was performed to determine whether within-subject differences in orientation depended upon corresponding differences in mean endpoint position. For this analysis, differences in orientation were calculated not as differences in azimuth and elevation but as differences in eigenvector orientation in space. Data from all directions were used.

#### Analyses between visual conditions

We also analyzed how differences in ellipsoid orientation between postures varied with movement extent and between visual conditions. For each visual condition, analyses were conducted at 0-100% of the total movement extent (in 5% increments). At each increment, repeated estimates of orientation difference were obtained by bootstrap resampling the hand position distributions associated with each arm posture 1000 times, calculating the orientations of the principal axes of variability and taking the difference. Two-factor circular ANOVAs were then conducted on these ellipsoid orientation differences using ‘visual condition’ and ‘movement direction’ as factors. To assess the variability inherent in these calculations we used a 2^nd^ bootstrap analysis. For each increment of movement extent, differences in orientation were calculated using two estimates of orientation obtained by bootstrap resampling hand position distributions associated with the *same* posture and visual condition. This bootstrap procedure was repeated 1000 times to obtain a distribution of orientation differences for each visual condition and each increment in movement extent.

## Results

As expected, initial arm orientations differed between the two instructed arm postures but were consistent between visual conditions and across movement directions (Table I). For the ADD posture, average arm orientations (rounded to the nearest degree) were approximately 60° for all movement directions in both conditions. For the ABD posture, average arm orientations were approximately 25° for all directions in both conditions. The ~35° difference in initial arm posture did not result in appreciable changes in standard behavioral or kinematic performance measures. When data for all targets were combined, mean reaction times were slightly longer for the ADD posture (10 msec longer for V; 19 msec for NV). However, these differences represented only a 4 and 5% increase in reaction times for the V and NV conditions, respectively. Similarly, peak velocities were slightly greater for the ADD posture, which was largely due to an increase in movement times. When data were combined across targets, movement times were slightly shorter for the ADD posture, though the average difference for all targets (25 msec for V; NV: 35 msec) represented only a 3 and 4% decrease in movement times for the V and NV conditions, respectively.

**Table I.**

| Visual conditions |  | V |  |  |  |  |  | NV |  |  |  |  |
| --- | --- | --- | --- | --- | --- | --- | --- | --- | --- | --- | --- | --- |
| Movement direction |  | L |  | M |  | R |  | L |  | M |  | R |
| Initial posture (degrees) | ADD | 60 +/- 5 |  | 59 +/- 5 |  | 59 +/- 5 |  | 60 +/- 5 |  | 60 +/- 5 |  | 60 +/- 5 |
|  | ABD | 25 +/- 5 |  | 24 +/- 5 |  | 25 +/- 5 |  | 25 +/- 5 |  | 24 +/- 5 |  | 25 +/- 5 |
| Peak Velocity (cm/s) | ADD | 36.8 +/- 6.9 |  | 35.2 +/- 6.3 |  | 32.5 +/- 6.2 |  | 36.7 +/- 8.2 |  | 34.2 +/- 6.7 |  | 33.1 +/- 6.6 |
|  | ABD | 33.3 +/- 5.2 |  | 32.4 +/- 5.6 |  | 29.9 +/- 4.0 |  | 34.6 +/- 7.9 |  | 32.8 +/- 7.2 |  | 30.8 +/- 6.6 |
| Movement time (sec) | ADD | 0.699 +/- 0.12 |  | 0.743 +/- 0.10 |  | 0.776 +/- 0.10 |  | 0.715 +/- 0.13 |  | 0.750 +/- 0.10 |  | 0.776 +/- 0.12 |
|  | ABD | 0.717 +/- 0.89 |  | 0.771 +/- 0.12 |  | 0.804 +/- 0.12 |  | 0.735 +/- 0.15 |  | 0.788 +/- 0.11 |  | 0.823 +/- 0.12 |
| Path length (cm) | ADD | 11.9 +/- 1.4 |  | 12.3 +/- 1.4 |  | 12.0 +/- 1.7 |  | 13.1 +/- 1.5 |  | 12.8 +/- 1.7 |  | 12.7 +/- 1.6 |
|  | ABD | 11.6 +/- 1.3 |  | 11.8 +/- 1.4 |  | 11.5 +/- 1.2 |  | 12.6 +/- 1.4 |  | 12.4 +/- 1.6 |  | 12.3 +/- 1.7 |
| Reaction time (sec) | ADD | 0.388 +/- 0.58 |  | 0.396 +/- 0.47 |  | 0.380 +/- 0.52 |  | 0.387 +/- 0.66 |  | 0.377 +/- 0.57 |  | 0.363 +/- 0.42 |
|  | ABD | 0.377 +/- 0.35 |  | 0.375 +/- 0.42 |  | 0.382 +/- 0.27 |  | 0.356 +/- 0.48 |  | 0.362 +/- 0.53 |  | 0.353 +/- 0.42 |

Figure 3 shows horizontal plane views of the movement endpoints and variability ellipsoids for a single subject in each experimental condition. In the V condition, the spatial distributions of the endpoints were relatively compact, resulting in smaller ellipsoids. In contrast, endpoints in the NV condition were more dispersed, indicating that the absence of hand visual feedback led to greater overall variability. Ellipsoids in the V condition were also more anisotropic and more consistent in orientation for a given movement direction than those in the NV condition. These effects are consistent with previous work showing that vision generally reduced the spread and altered the spatial distribution of movement endpoints during performance of both memory-guided and reaction time tasks (McIntyre et al., 1997;1998;Carrozzo et al., 1999;Apker et al., 2010;Apker and Buneo, 2012). For this subject, initial arm posture appeared to have negligible effects on ellipsoid size, shape and orientation in the V condition. In the NV condition however, changes in posture resulted in noticeable differences in ellipsoid orientation, but these rotations were not consistent in sign or magnitude across movement directions.

**Figure 3.**
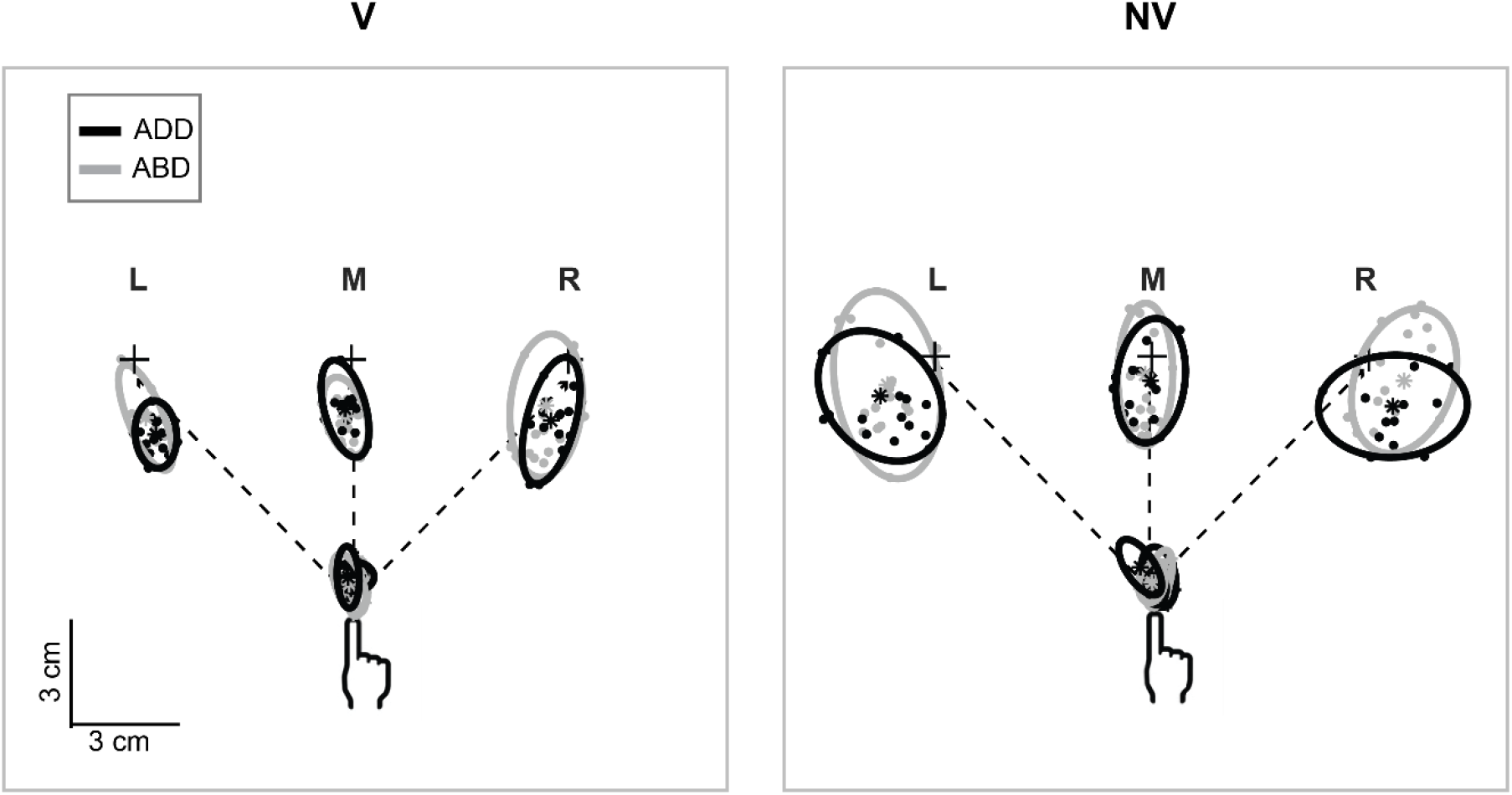
Top-down view of movement starting points and endpoints and associated 95% confidence ellipses for reaches to the left (L), middle (M) and right (R) targets for a representative subject. Dashed lines show the straight line paths to each target.

Figure 4 shows a sagittal view of the movement endpoints and variability ellipsoids from the same subject. As in the horizontal plane, movement endpoint distributions appeared more compact in the V condition compared to the NV condition. Ellipsoid orientations were also more consistent in the V condition. Interestingly, axes of maximum variability were not well aligned with planned hand movement directions (dashed lines). Instead, these axes were better aligned with the approximate lines of sight, suggesting variability was more strongly influenced by uncertainty in visually estimating the position of the hand and/or targets in depth than by execution related factors. In the NV condition, ellipsoid sizes, shapes and orientations were more variable and showed no particular alignment to planned movement directions or lines of sight. Arm posture had little effect on ellipsoid sizes and orientations in the V condition and again showed inconsistent effects in the NV condition. In summary, in this subject the V condition was associated with variability ellipsoids that were relatively small and consistent in orientation, while in the NV condition ellipsoids were larger and more variable in orientation across directions and between arm postures.

**Figure 4.**
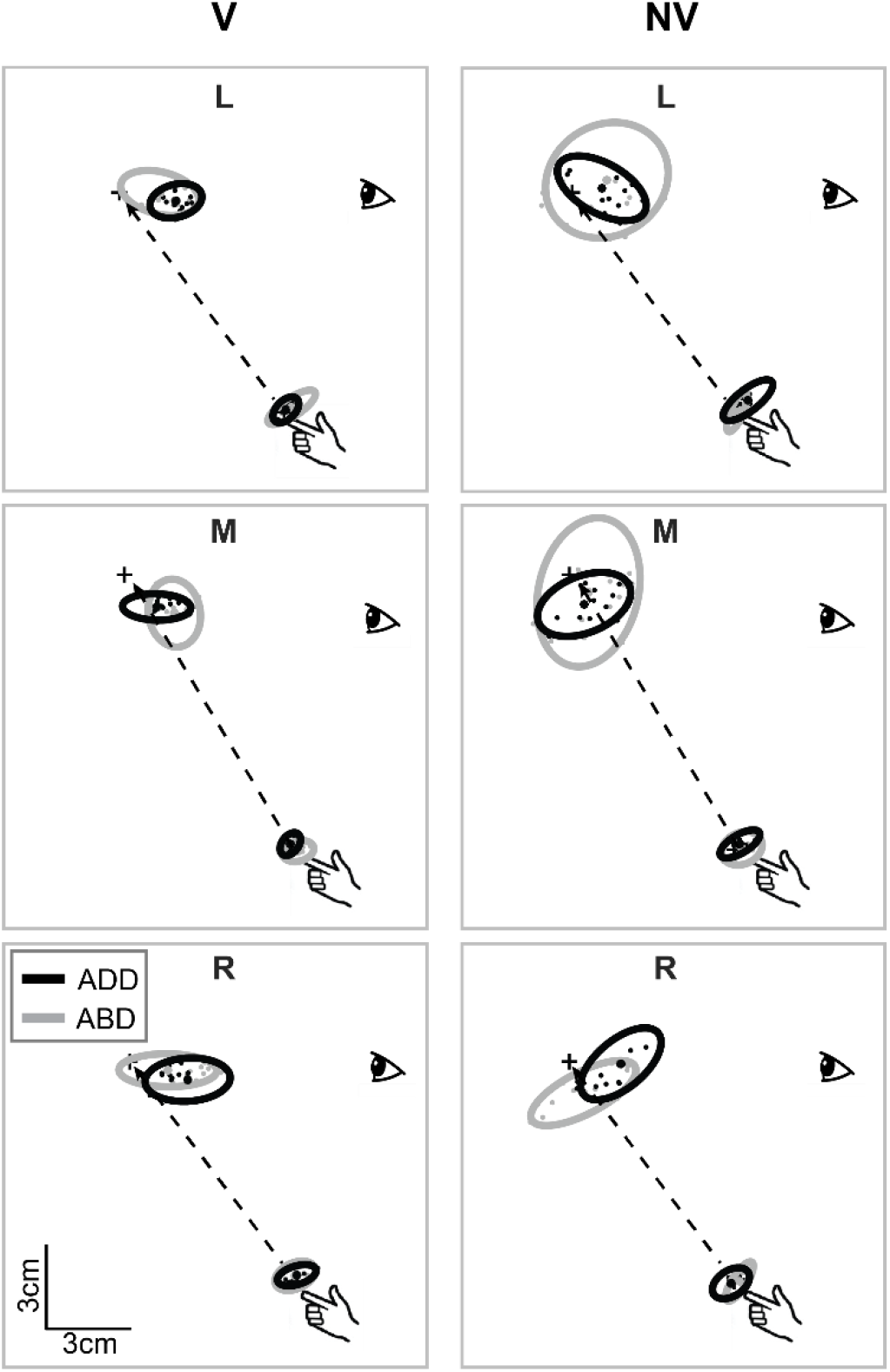
Lateral view of movement endpoints and confidence ellipses for reaches to each target for the subject shown in Fig. 3. Dashed lines show the straight line paths to each target.

Systematic effects of arm posture on ellipsoid size and shape were also not consistently evident at the population level. Figure 5 shows bar plots of the ellipsoid volumes and aspect ratios for all experimental conditions. In the V condition (left column), average volumes were generally small and varied little across movement directions. In the NV case, volumes were typically more than twice as large, but still varied little with movement direction. Although volumes were also generally much more variable across subjects in this condition, more striking was the lack of consistency in how volume varied between postures from subject to subject (light grey points and lines). For a given direction, these differences were often large and opposite in sign between subjects. ANOVA confirmed these observations: for the V condition there was a significant main effect of posture (*F*(1,54) = 6.00, *p*<0.05) but no significant effect of movement direction (*F*(2,54) = 0.38, *p*=0.962) and no significant interaction between movement direction and posture (*F*(2,54) = 0.52, *p*=0.607). For the NV condition, no effects of movement direction (*F*(2,54) = 1.49, *p*=0.264) or posture (*F*(1,54) = 0.21, *p*=0.667) were found and there was no significant interaction effect (*F*(2,54) = 0.09, *p*=0.914).

**Figure 5.**
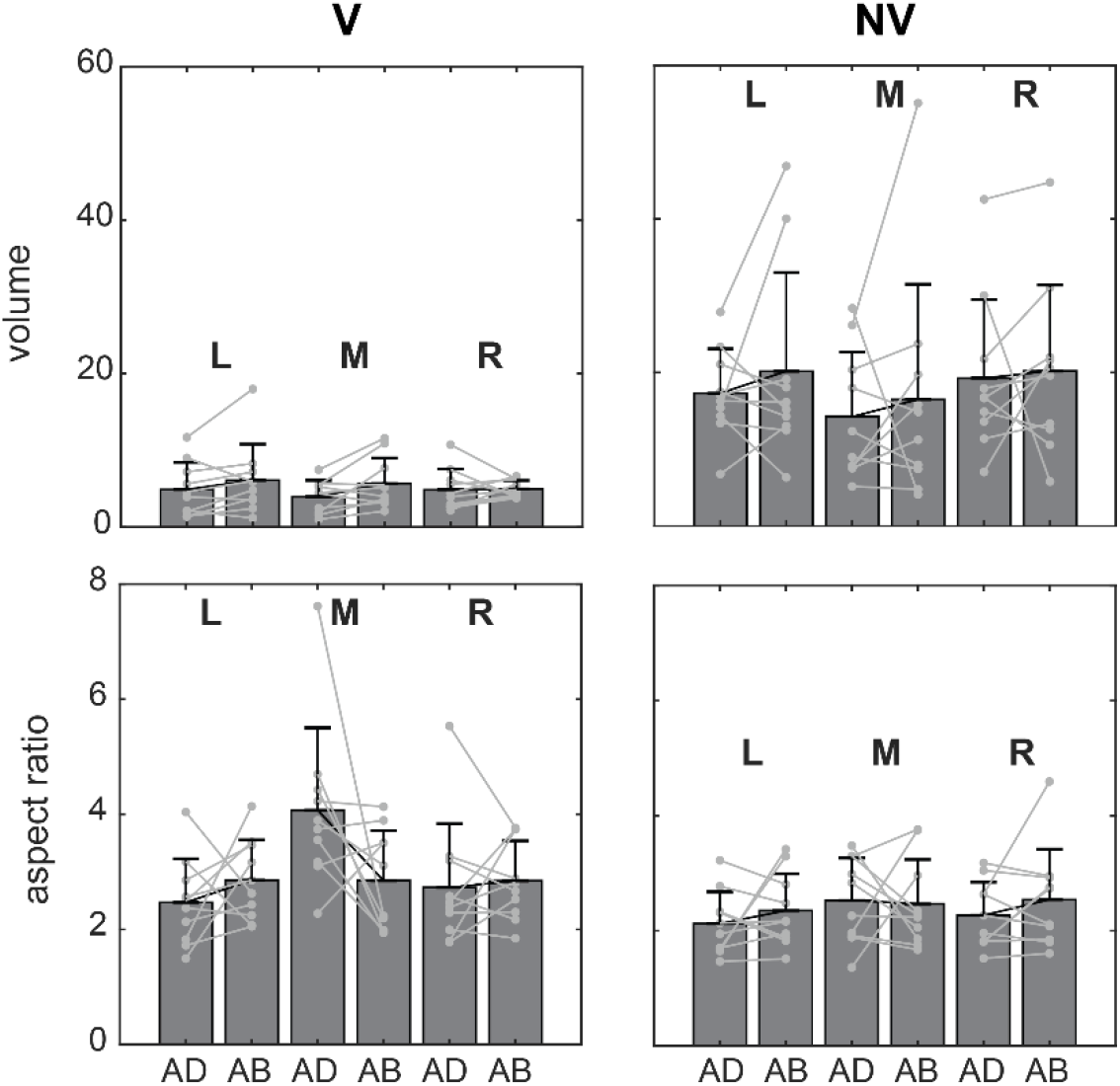
Average volumes and aspect ratios (+/− SD) for reaches to each target. Individual subject data are superimposed on each set of bars (light gray lines). Average volumes were consistently larger in the NV condition but showed inconsistent variation with target direction and arm posture. Average aspect ratios were more similar between vision conditions and arm postures as well as across targets.

Regarding ellipsoid shapes, aspect ratios in the V condition were somewhat larger than in the NV condition for all movement directions. Larger aspect ratios indicate more elongated ellipsoids, which were also apparent in the V condition for the example subject in Figs. 3 & 4. In both visual conditions, effects of arm posture on aspect ratio were observable but did not follow any systematic trend. These observations were borne out by ANOVA which showed that in the V condition there was a significant effect of movement direction on aspect ratio (*F*(2,54) = 3.82, *p*<0.05) but no main effect of posture (*F*(1,54) = 0.05, *p*=0.830) and no significant interaction between direction and posture (*F*(2,54) = 2.67, *p*=0.104). For the NV condition no main effects of movement direction (*F*(2,54) = 0.68, *p*=0.517) or posture (*F*(1,54) = 1.123, *p*=0.297) were found and there was no significant interaction effect (*F*(2,54) = 0.340, *p*=0.716). Overall, the data in Fig. 5 suggest that vision had a larger effect on volume than aspect ratio and that arm posture had strong effects on both ellipsoid parameters (particularly in the NV condition), but due to their idiosyncratic nature, these effects were not captured by the statistical analyses.

At the population level, effects of arm posture on ellipsoid *orientation* were also inconsistent across subjects. Figure 6 shows horizontal plane views of the principal axes of variability for all subjects in each condition, which allows visualization of the azimuth component of orientation. Black lines represent the group average and grey lines represent individual subjects. In the V condition, individual axes were oriented closer to the average and were less dispersed than those in the NV condition. In addition, looking across directions, average axes were more convergent toward the starting hand position (and therefore body midline) in the V condition than in the NV condition. In the V condition, effects of arm posture were not obvious when data were viewed in this plane. In the NV condition, axes for movements to the leftward target appeared to rotate in a clockwise manner as initial arm posture was varied from ADD to ABD. Otherwise, for both ADD and ABD orientations appeared very similar. Therefore, although visual feedback resulted in more spatially convergent axes of variability, arm posture changes appeared to have little additional effect on the azimuthal orientation of these axes.

**Figure 6.**
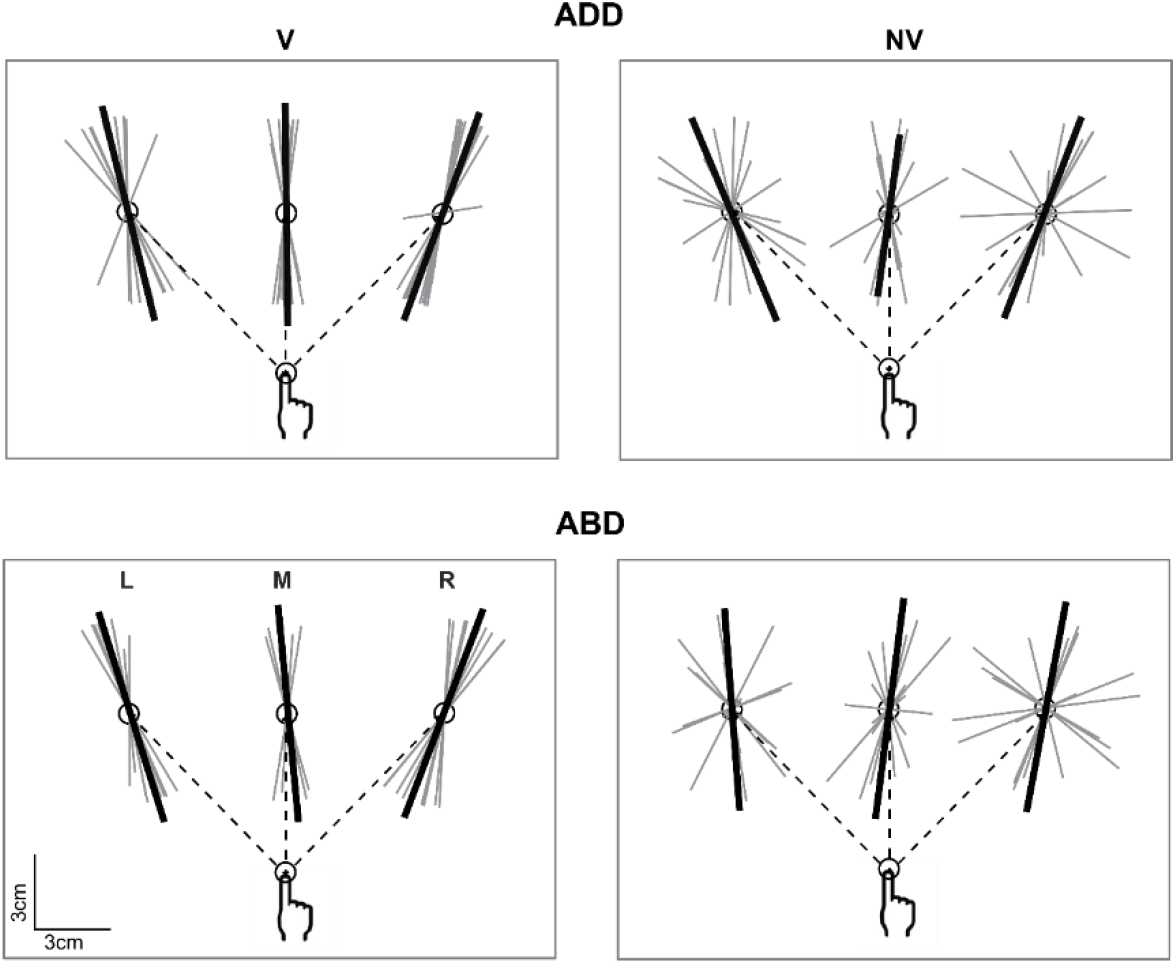
Top-down view of the individual (gray) and average (black) principle axes of variability in the V condition. Axes were scaled to an arbitrary length for visualization purposes. Dashed lines show the straight line paths to each target. For both arm postures, individual vectors were generally less variable in orientation and average vectors were more convergent across targets than in the NV condition.

Larger effects of arm posture were evident when eigenvector orientation was examined in the sagittal plane. In the V case (Fig. 7), average axes showed little variation in elevation angle across directions or between arm postures. Axes were generally aligned along subjects’ sight lines to the targets, as was shown for the individual subject in Fig. 4. In contrast, individual axes in the NV condition varied widely across directions and between postures and average axes were inconsistently oriented (Fig. 8). For example, for some directions (L for ADD, M for ABD) average axes aligned approximately with the sight lines. For others (M for ADD, L and R for ABD) these axes were either pitched strongly upward, reminiscent of previous findings for memory-guided reaches (McIntyre et al., 1997;1998), or strongly downward (R for ADD).

**Figure 7.**
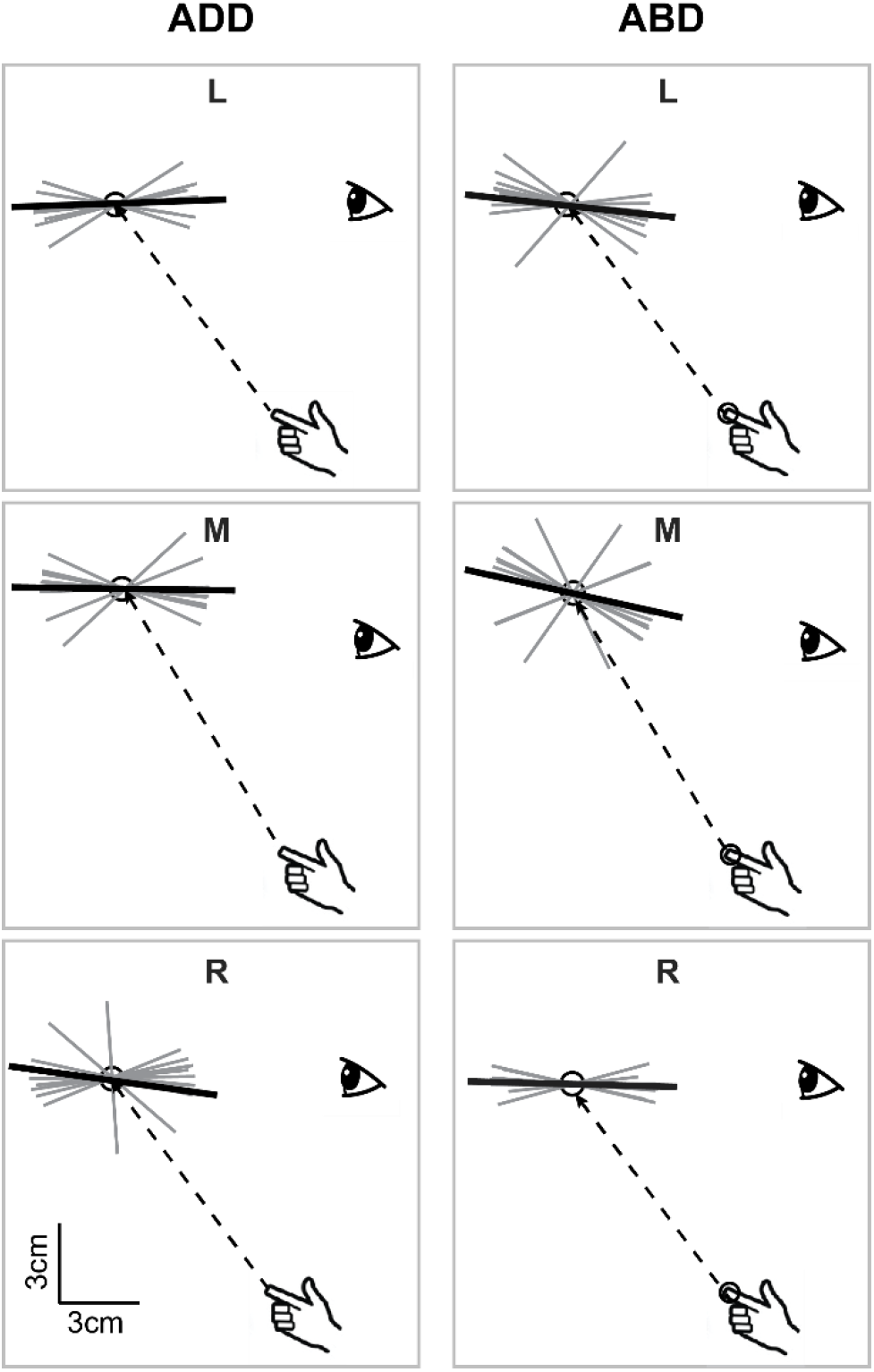
Lateral view of the individual (gray) and average (black) principle axes of variability in the V condition. For both arm postures, average vectors were generally collinear with the sight line to the target and were not well-aligned with planned movement vectors (dashed lines).

**Figure 8.**
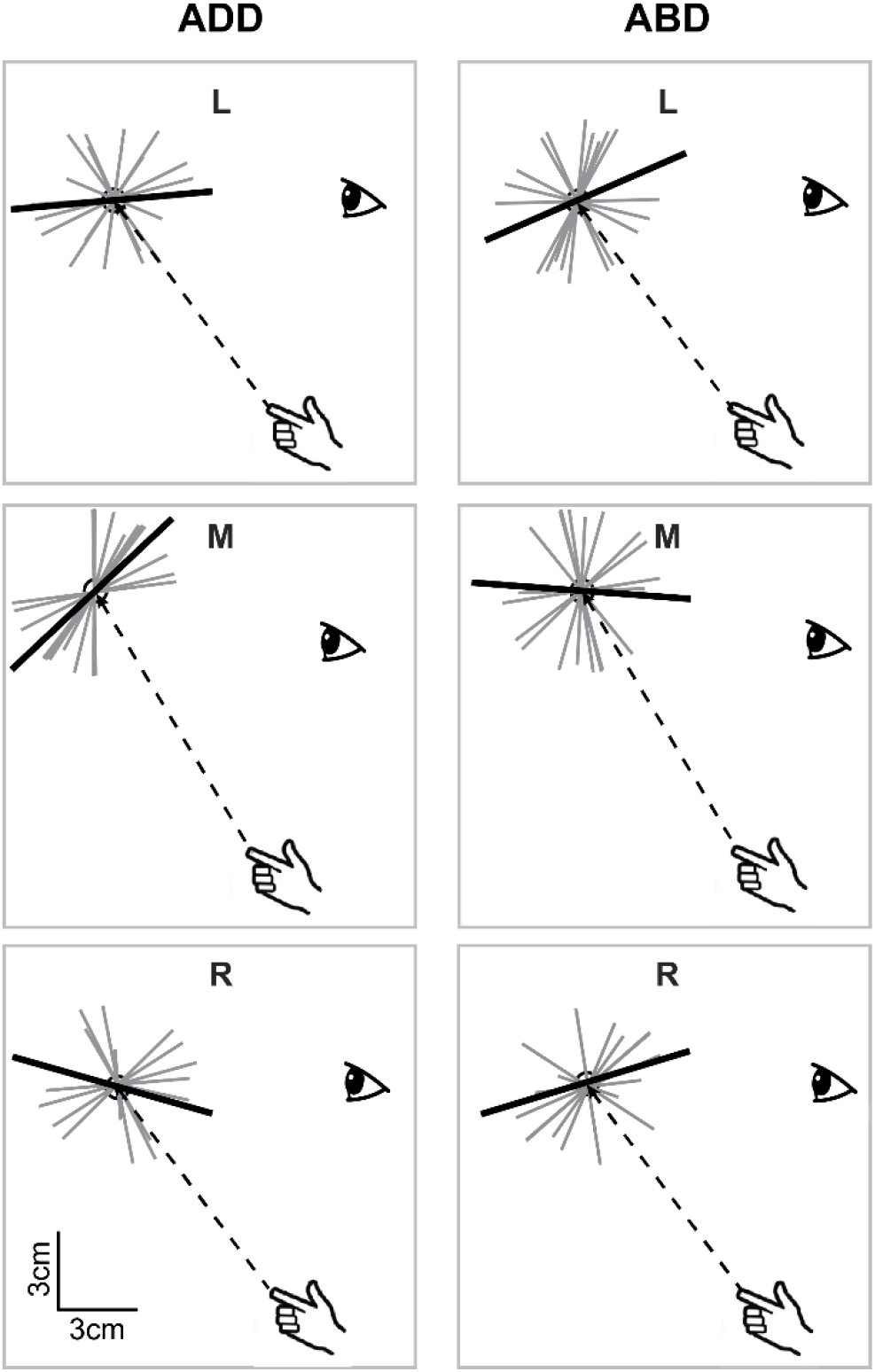
Lateral view of the individual and average principle axes of variability in the NV condition. For both arm postures, individual vectors were generally more variable in orientation than in the V condition. In addition, average vectors were generally more divergent across targets and between postures than in the V condition.

Although average orientations appeared to vary somewhat randomly with direction and arm posture in Figs. 6–8, it was difficult to appreciate from these plots how such changes manifested at the single-subject level. Figure 9 shows the ellipsoid orientation data for single subjects superimposed on bar plots of the average orientations. For the V condition, average azimuth angles grossly followed changes in the required movement direction, decreasing as movement direction varied from left to right. Average elevation angles showed less variation, hovering around 0°, indicative of largely horizontal orientations. Arm posture appeared to have little consistent effect on these average angles. In the NV condition, the situation was very different. Here, mean azimuth angles still grossly followed changes in movement direction for both arm postures but variability was markedly increased relative to the V condition. Importantly, the direction in which azimuth changed with arm posture was again quite variable across subjects, with some showing large increases in azimuth from AD to AB and others showing the opposite trend. Mean elevation angles were even more variable across directions and, similar to the azimuthal data, varied widely between postures for the same subject.

**Figure 9.**
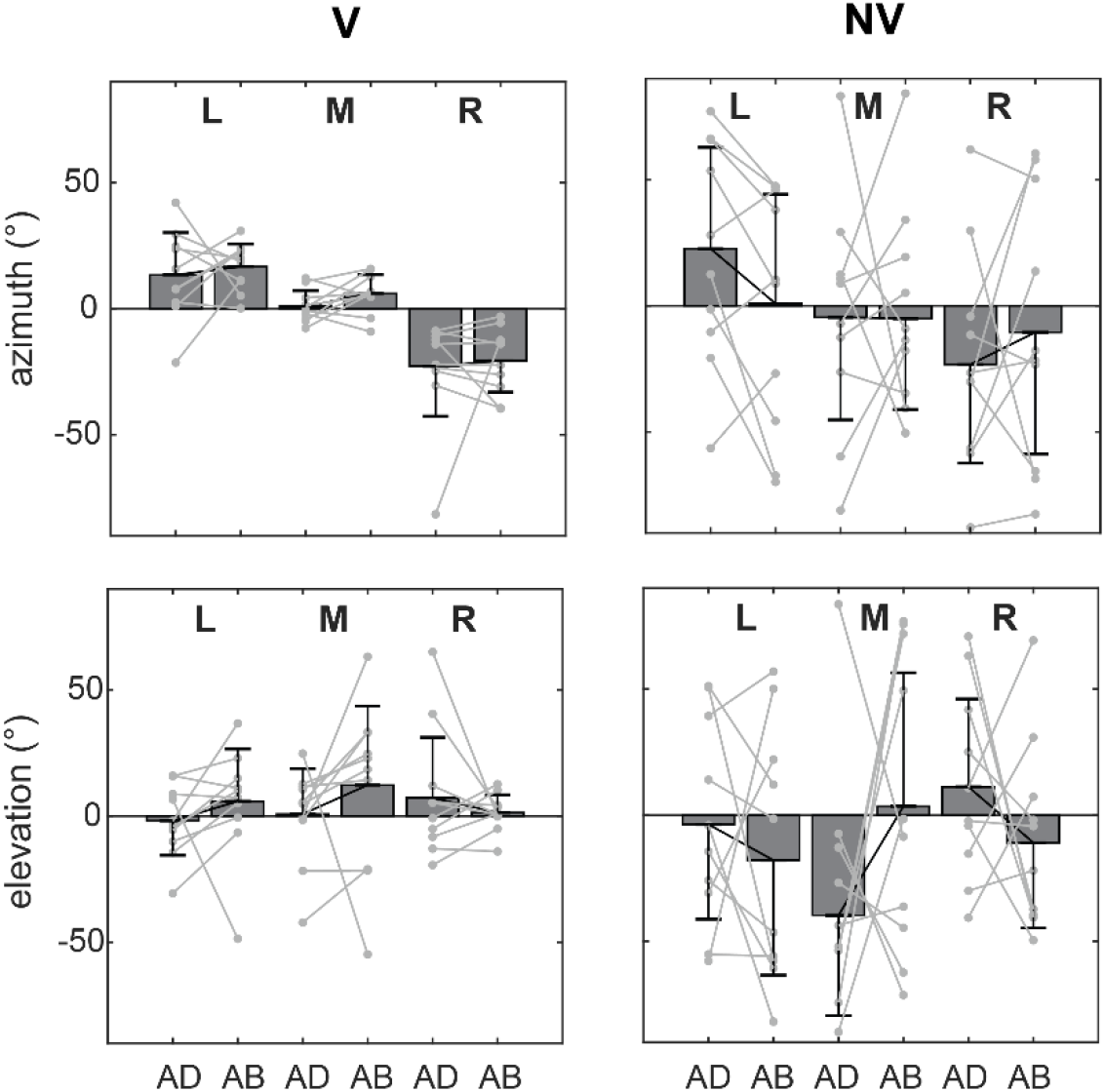
Average orientations of the principle axes of variability (+/− circular SDs) for each target. Individual subject data are superimposed on each set of bars (light gray lines). Average values of azimuth varied systematically with target direction in the V condition and to a lesser extent in the NV condition. Average elevation values showed less consistent variation. Both azimuth and elevation were markedly more variable in the NV condition and trends with arm posture were highly idiosyncratic.

These qualitative observations of the orientation data were largely confirmed by statistical analysis. In the V condition there was a statistically significant effect of movement direction on azimuth (*F*(2,54) = 36.60, *p*<0.05) but no significant effect of posture (*F*(1,54) = 0.97, *p*=0.329) and no significant interaction between movement direction and posture (*F*(2,54) = 0.17, *p*=0.840). For the NV condition, no effects of movement direction (*F*(2,54) = 1.61, *p*=0.209) or posture (*F*(1,54) = 0.04, *p*=0.838) on azimuth were found and there were no significant interaction effects (*F*(2,54) = 0.58, *p*=0.563). Regarding elevation, there were no significant effects of movement direction, posture, or their interaction in either the V condition (*F*(2,54) = 0.34, *p*=0.716; *F*(1,54) = 0.60, *p*=0.443; *F*(2,54) =1.12, *p*=0.337) or the NV condition (*F*(2,54) = 1.20, *p*=0.309; *F*(1,54) = 0.08, *p*=0.783; *F*(2,54) =2.06, *p*=0.137). Although these analyses pointed to a lack of postural effects, Fig. 9 shows that this resulted from a lack of *systematic* variation with arm posture. Importantly, strong postural effects were clearly evident in the data in this figure but were idiosyncratic in nature.

The data in Fig. 9 suggest that changes in variability ellipsoid orientation due to arm posture were not systematic across subjects but idiosyncratic, changing in both magnitude and direction in a subject specific manner. In addition, this figure suggested that these idiosyncratic changes were much greater in the NV condition. To quantify these effects we performed a circular ANOVA on the within-subject differences in orientation between the ADD and ABD postures, using movement direction and visual condition as factors. We found a statistically significant effect of visual condition on these within-subject differences (*F*(1,54) = 23.54, *p*<0.05), but no effect of movement direction (*F*(2,54) = 0.43, *p*<0.653) and no interaction between movement direction and vision condition (*F*(2,54) = 0.06, *p*<0.941). Thus, within-subject, posture-related differences in ellipsoid orientation were strongly dependent upon the presence/absence of visual feedback and were independent of movement direction.

Previous simulation studies predicted strong, systematic effects of movement direction and arm posture on movement variability that were not observed here. However, these simulations did not incorporate feedback and consequently analysis was focused on variability in initial movement direction, not endpoint variability. Although we observed idiosyncratic effects of posture on endpoint variability in the present study, it is possible that at earlier points in the movements effects were more systematic, as our simulations predicted. To explore this possibility, we calculated the average within-subject differences in orientation between arm postures as a function of movement extent for both feedback conditions (Fig. 10). This analysis showed that differences in orientation were observed from the very beginning of movement. These differences were not systematic however, as no statistically significant effects of arm posture on orientation (azimuth/elevation) were observed at any point during the movement. Interestingly, differences between visual conditions did not emerge until very late, i.e. at approximately 80% of the total movement extent. At that point, orientation differences between the two arm postures increased markedly in the NV condition but did not in the V condition. However, a bootstrap analysis (inset) showed that most of the observed differences in orientation in the V condition could be explained by the variability inherent in the calculation of orientation differences. In contrast, difference in the NV condition that emerged late in the movement were substantially larger than what one would expect based on this inherent variability.

**Figure 10.**
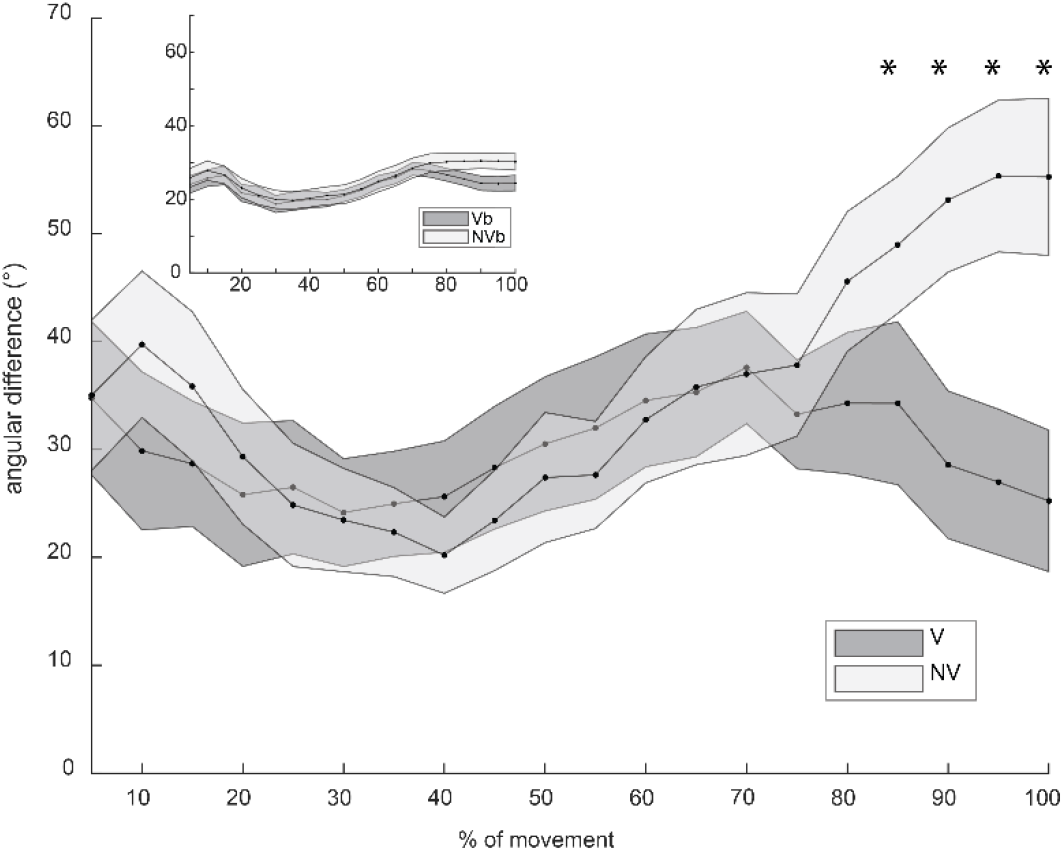
Average within-subject differences in orientation between arm postures, plotted as a function of movement extent for both feedback conditions. Error bars represent standard deviations. Inset: bootstrap analysis showing expected differences in orientation when using data from the same posture and condition. See text for details.

Our analyses showed that idiosyncratic differences in variability due to initial arm posture were present from the very beginning of movement but became magnified toward the end of movement when vision of the moving hand was unavailable. What could give rise to these differences? One possibility is that they are related in some way to differences in final endpoint position between conditions. If this were the case it could implicate final limb impedance, which is position dependent (Mussa-Ivaldi et al., 1985;Artemiadis et al., 2010) and has been shown to influence movement variability for planar reaching movements (Lametti and Ostry, 2010), as a factor in determining the orientation of endpoint variability ellipsoids in this experiment. To explore this possibility we regressed within-subject differences in ellipsoid orientation on within-subject differences in final position for all targets and subjects. Figure 11 shows scatterplots of orientation difference vs position difference for both conditions. These variables were uncorrelated in the V condition but were moderately positively correlated in the NV condition. Moreover, a linear regression analysis showed that in the V condition, differences in final position did not account for a significant proportion of the variance in orientation difference (p=0.62; F= 0.25; R^2^= 0.01). In the NV condition however this regression was statistically significant, accounting for approximately 15-20% of the variance in orientation difference (p=0.03; F= 5.32; R^2^= 0.17). Thus, in the NV condition differences in the final orientation of variability ellipsoids in this study depended in part on differences in final endpoint positions.

**Figure 11.**
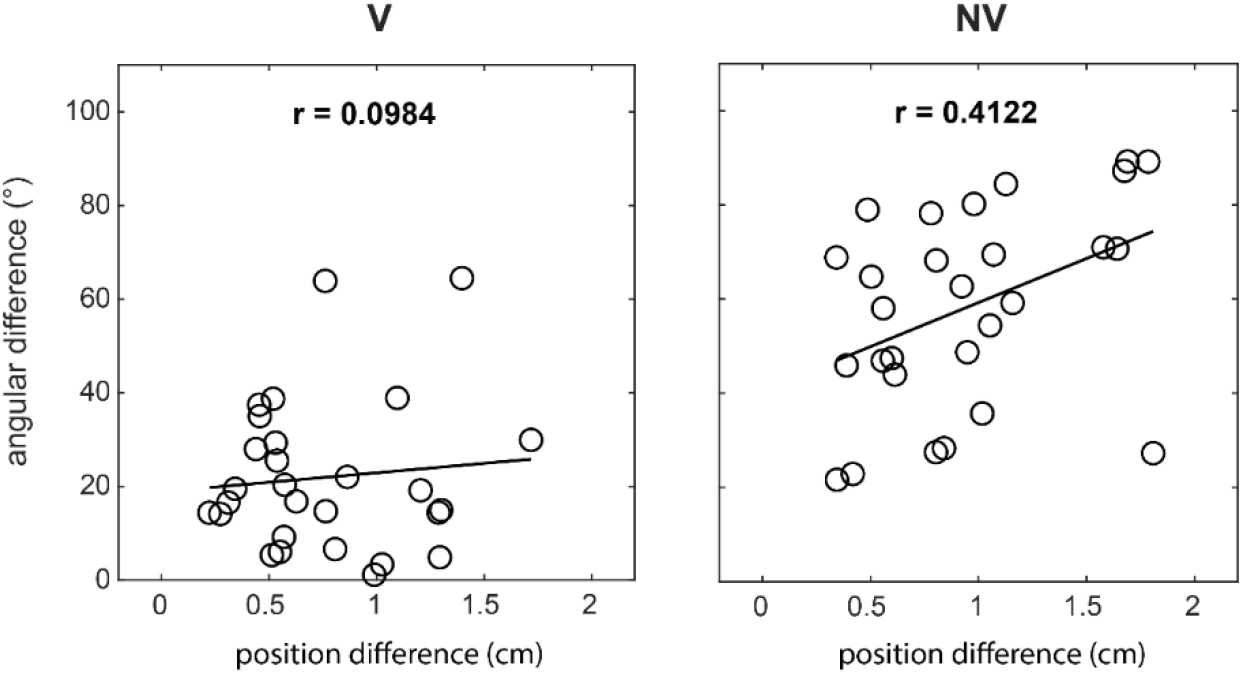
Scatterplots of average orientation difference vs average position difference for both conditions. Data from all targets and subjects are shown.

## Discussion

Here we examined the effects of changes in initial arm posture on movement variability in 3D space. Subjects executed memory-guided reaching movements in three directions using one of two initial arm configurations, which were attained by rotation about the shoulder-hand axis. In this way, effects of arm posture were examined for nominally identical sets of planned endpoint trajectories. In addition, movements were performed with and without vision of the moving hand (V and NV conditions, respectively). We found that reach endpoint distributions differed in orientation between initial arm configurations, but in a subject-dependent way. These differences were largest during the terminal phases of movement for reaches made in the NV condition. Moreover, in this condition, within-subject differences in mean endpoint position were moderately predictive of corresponding differences in the average orientations of reach endpoint distributions. As discussed below, the results emphasize the role of factors such as biomechanics and suboptimal sensorimotor integration (rather than internal noise) in constraining patterns of movement variability in 3D space.

### Relation to previous psychophysical studies

Several previous studies have examined the contributions of planning and execution-related noise to movement variability for both planar (2D) arm movements (van Beers et al., 2004; Gordon et al., 1994; Vindras et al., 1998) and reaching or other arm-related behaviors in 3D space (Apker, 2010, 2012; Carrozzo et al., 1999; McIntyre, 1997, 1998; van den Dobbelsteen et al., 2001). The results of the 3D reaching studies are most germane to the present one. In the V condition we found a significant effect of movement direction on the orientation of axes of maximum variability. Moreover, these orientations appeared to vary systematically with the direction of the targets relative to the head and/or eyes, rather than with respect to the starting position of the hand. This suggests that the effect of direction in this condition is strongly influenced by noise in the initial planning (and/or updating) of hand position in viewer/eye-centered coordinates, rather than noise in hand/arm-centered coordinates, consistent with the conclusions of McIntyre and colleagues (McIntyre et al., 1997;1998). Effects of arm posture on orientation in this condition were largely idiosyncratic (Fig. 9). Moreover, when differences in orientation between arm postures were analyzed as a function of movement extent, they were found to be relatively constant and small (~10°or less). Thus, the present results reinforce previous findings implicating visually derived planning noise as a strong determinant of endpoint variability when vision of the hand is available, with arm posture influencing variability only slightly and in a subject-dependent way.

In the NV condition, variability was larger, as expected, and there was no significant effect of direction, arm posture or the interaction of these factors on the absolute orientations of the endpoint distributions. Figures 6 and 8 showed that endpoint distributions were less convergent toward the head in this condition and also did not align well with hand movement direction, findings that were also consistent with those of McIntyre and colleagues. Similar to the visual condition, but more dramatically, Fig. 9 showed that orientation varied greatly between postures in both magnitude and direction, but in a manner that was again largely idiosyncratic to each subject. When relative changes in orientation were analyzed as a function of movement extent, they did not differ from those in the V condition early in the movement but grew much greater in the terminal phases. After accounting for inherent variability in the calculation of orientation differences within each condition, orientations were found to differ by ~25° by the end of the movement, much greater than in the V condition.

What conclusions can be drawn from the orientation data in the NV condition? First, the observation of large differences in orientation between postures (even if idiosyncratic) indicates that previously described differences in orientation when movements were initiated with different hands from the same starting position (McIntyre et al., 1998) were not due entirely to effects of handedness, but were at least partly posture dependent. This is also supported by recent results from a reaction time task, which showed that handedness was associated only with differences in the overall size and not the shape or orientations of movement endpoint distributions (Apker et al., 2015). Second, the lack of a consistent pattern of orientation change with direction relative to the head/eyes or the starting hand position suggests that endpoint variability in this condition cannot be explained by planning noise in visual coordinates or execution noise in hand/arm coordinates. An alternative explanation is that the observed patterns of variability resulted from suboptimal sensorimotor integration. That is, a recent report showed that the proprioceptive map of the arm is stable but non-uniform in structure across subjects (Rincon-Gonzalez et al., 2011). This non-uniformity may result from idiosyncratic neural approximations to the highly non-linear mapping between extrinsic (endpoint) and intrinsic (joint) space (Flanders et al., 1992). Simulations suggest that complex problems such as integrating an estimate of the hand’s position based on feedforward commands with an idiosyncratic approximation of the hand’s position derived from proprioceptive feedback can result in suboptimal estimation, leading to increased behavioral variability (Beck et al., 2012). It is conceivable therefore, that a similar process could have contributed to the large and gradually evolving differences in orientation between postures that was observed in the NV condition.

Work by other groups suggests that biomechanical factors, specifically limb impedance, can also explain some aspects of the posture dependent variability observed here. Importantly, limb impedance has previously been implicated as a factor influencing behavior (including variability) during planar arm movements (Scheidt and Ghez, 2007;Lametti and Ostry, 2010) and is arm position/configuration dependent in both 2D and 3D space (Mussa-Ivaldi et al., 1985;Artemiadis et al., 2010). Our regression analysis, which showed that orientation differences were correlated with differences in mean endpoint position in the NV condition, is consistent with the idea that position dependent differences in limb impedance contributed to the larger differences in variability observed in this condition. A stronger influence of final limb impedance on variability in the NV condition could help explain the observation that axes of maximum variability did not align well with the eyes/head or planned movement direction and did not depend in any consistent way on starting arm posture. It could also explain the discrepancy between the present experimental results and our previous simulations (as discussed below), as the latter did not incorporate a separate position controller.

### Relation to previous simulation studies

Based in part on the results of previous simulation studies (Shi and Buneo, 2012), we predicted that patterns of endpoint variability would rotate in a systematic manner with changes in arm posture, a prediction that was not observed. There are at least two possible factors contributing to this discrepancy. The first concerns the nature of the arm configuration changes in the present study, which involved rotations about the shoulder-hand axis. Such changes in posture are largely irrelevant to the planning of hand movement vectors (as they don’t change the position of the hand relative to the goal location) though they are still highly relevant to the planning of dynamics (Soechting et al., 1995;Buneo et al., 1997). In contrast, changes in arm posture that are coplanar with planned movements (as in Shi and Buneo (2012)) affect the computation of *both* movement vectors (through their effects on initial hand position) and dynamics (through their effects on elbow torque) (Buneo et al., 1995;Sober and Sabes, 2003). The combined result of these effects are patterns of variability that vary systematically with both direction and posture. Thus, differences in the relative contributions of movement vector planning noise between the simulation studies and the present experiments could be partially responsible for the observed discrepancy with regard to postural effects.

A second possible factor leading to the observed discrepancy between our previous simulations and the present experimental results concerns the role of feedback. That is, these simulations were entirely feedforward whereas in contrast, movements in the present study were closed loop with respect to proprioceptive feedback (in both conditions) and visual feedback in the V condition. Thus, it is possible that the presence of online feedback interacts with feedforward commands in such a way that effects of arm posture on movement variability appear less systematic than expected. However, it should be noted that idiosyncratic effects of arm posture were observed from the very beginning of the movement in both sensory conditions, i.e. well before online feedback could have influenced the ongoing motor command. Thus, it seems unlikely that differences in the feedback conditions were the sole factor leading to the non-systematic changes in variability observed here.

## Acknowledgments and Support

Supported by National Science Foundation Division of Integrative Organismal Systems Award 1558151

## Notes

### Competing Interest Statement

The authors have declared no competing interest.

